# Exploring the impact of interthalamic adhesion on human cognition: insights from healthy subjects and thalamic stroke patients

**DOI:** 10.1101/2024.02.08.579448

**Authors:** Julie P. Vidal, Kévin Rachita, Anaïs Servais, Patrice Péran, Jérémie Pariente, Fabrice Bonneville, Jean-François Albucher, Lola Danet, Emmanuel J. Barbeau

## Abstract

**Background and Objectives:** Interthalamic adhesion (IA), also known as the massa intermedia, is a structure that connects the median borders of both thalami across the third ventricle. Given it is difficult to identify on routine neuroimaging, its anatomical variants and function remain poorly studied. The main objective of this study was to clarify the role of IA on cognition. Our main hypothesis was that thalamus stroke patients *with* an IA would show better performance on neuropsychological tests than individuals *without* an IA through possible compensatory mechanisms.

**Methods:** The study comprised a group of healthy subjects and a cohort of patients with isolated thalamic strokes at the chronic stage. All participants underwent 3T research T1w and FLAIR MRI as well as a neuropsychological assessment. The presence or absence of an IA and type of IA anatomical variant were evaluated by two independent reviewers.

**Results:** 42 healthy subjects (mean age= 49) and 40 patients (mean age= 51) were finally included. 76% of participants had an IA, with a higher prevalence among women (92%) than men (61%). The presence or absence of an IA did not effect the neuropsychological performance of healthy subjects nor did the type of IA variant. Across all the tests, patients *with* an IA (n = 18) showed the lowest BF_10_ (157) while those *without* an IA (n = 10) exhibited the highest BF_10_ (10648) when compared to healthy subjects using a Bayesian rmANOVA. More specifically, patients *without* an IA performed more poorly in the verbal memory or Stroop task versus healthy subjects than patients *with* an IA. This effect was not explained by age, laterality of the infarct, volume or, localization of the lesion. Patients with an IA and lesions extending into the IA presented a similar trend to non-IA subjects which could however be explained by a greater volume of lesions.

**Discussion:** IA does not appear to have a major role in cognition for healthy subjects but could play a compensatory part in patients with thalamic lesions.

## 1. Introduction

Interthalamic adhesion (IA), or massa intermedia, is a midline structure connecting the median border of both thalami across the third ventricle, posterior to the foramen of Monro [1–4]. Intriguingly, the prevalence of IA within the population varies across MRI studies, ranging from 68% to 98%, with most studies reporting an absence of IA in around 20% of cases [2, 4, 12–16]. Its typical form in humans is a single and homogenous adherence usually located at the anterosuperior quadrant [1–4]. IA has anatomical variants: a broad, larger, form in 18% of cases and duplication which occurs in 2-10% of instances [1, 2, 4–9]. Other rare variants include bilobar, multiple, tubular or rudimental IA seen in less than 3% of subjects [8]. The “kissing thalami” phenomenon occurs when both thalami adhere to one another, preventing proper assessment of whether an IA is present or not [6, 10, 11]. Prevalence of IA is thought to be more common in females (91%) compared to males (83%) [4, 7, 10, 13, 14, 16, 17]. A larger IA is also reported among females versus males [1, 2, 8, 18]. Most studies used T1-weighted MRI scans with a great disparity among methods in the absence of a standardized protocol to evaluate prevalence, anatomical variants or size of IA.

The reasons for the absence of an IA are debated and include a potential correlation with age [6, 15, 18–21], third ventricle abnormalities [3, 5, 6, 13, 15, 20] and could possibly be related to various neurodevelopmental or psychiatric disorders such as schizophrenia, borderline personality, major depression or bipolar disorder [6, 16, 17, 20, 22–24]. Absence of an IA could be linked to early neurodevelopmental abnormalities during gestation [16, 21, 24].

Some studies found neuronal cell bodies in IA using Nissl-stained material [12] or Golgi material [25] while a recent histological study using hematoxylin-eosin staining pointed towards their absence [26]. Although the debate continues on whether IA should be considered as gray or white commissure, the recent discovery of axons in its core appears to provide evidence in favor of the latter [7, 27].

Due to its limited size, little is known regarding the connectivity supported by IA. Damle et al. [28] found that the IA size is associated with anterior thalamic radiations, related to dorsomedian thalamic nuclei and involved in memory [29]. Recent diffusion tensor imaging studies have identified crossing stria medullaris fibers passing through the IA, connecting the prefrontal cortex to the contralateral habenula via the internal capsule and anterior nuclei of the thalamus. When IA was too small, no crossing fibers were identified and in the absence of IA, fibers crossed via the posterior commissure [30]. IA has also been shown to be connected to the amygdala, hippocampus, temporal lobes, insular and pericalcarine cortices [11, 7].

The role of IA in human cognition has been a subject of uncertainty for a long time, with some authors considering it a vestigial part of the brain with no known function [9]. As already mentioned, the fact that IA is not present in all subjects probably reinforced the idea that it may not play a critical part in cognition. However, a few studies have started to shed light on its potential cognitive significance. The size of the IA was thought to mediate the relationship between age and attention in healthy female subjects [28]. A recent study using healthy subject data from the Human Connectome Project associated IA absence with inhibition and attention deficits as well as increased negative emotional function [10]. In addition, the absence of IA among patients with epilepsy was linked to worse performances on verbal memory tests and executive functions [18]. However, as mentioned by the authors, memory deficits in medial temporal lobe epilepsy patients can also be directly linked to epilepsy. Overall, the role of IA in human cognition has rarely been investigated using dedicated protocols. This may be due to several factors, including the need for high-quality MRI which is only recently available but necessary to identify IA and the difficulty of taking into account IA variants.

In this context, the aim of our study is to clarify the role of IA in human cognition using a novel approach as assessed in both a group of 45 healthy control subjects and 40 patients with isolated ischemic thalamic stroke. Our general hypothesis was that IA plays a part in cognition. More specifically, we thought that thalamic stroke patients might reveal the role of IA in cognition through functional compensation mechanisms. We hypothesized that patients *with* an IA would produce better performance than patients *without* an IA on neuropsychological tests. We did not make any specific hypothesis regarding the case of patients *with* an IA *and* a thalamic lesion encompassing their IA.

## 2. Materials and Methods

### 2.1 Participants

We collected data from 45 healthy subjects (ages 23-69, median age 52, 20 males) and 40 patients (ages 23-75, median age 54, 25 males) with ischemic thalamic lesions at the chronic stage (91-2,674 days post-stroke). These patients were recruited in the Stroke Units at University Hospital of Toulouse between 2011-2013 and 2019-2020.

Patients from two studies were included in order to increase the sample size. The first study was approved by the Institutional Review Board “Comité de Protection des Personnes Sud-Ouest et Outre-Mer no. 2-11-0”. It included 20 patients with a thalamic stroke, younger than 80 years old. Marginal extra-thalamic lesions were accepted in this study (described in [31]). The second study was authorized by the “Comité de Protection des Personnes Ile-de-France IV” (Ethics Committee). It included 20 patients aged below 70 years with at least one stroke lesion visually reaching the dorsomedian nucleus and no extra-thalamic damage. For both studies, recruitment criteria were detection of a first symptomatic thalamic infarct regardless of complaint or neurobehavioral report before onset and no previously known neurovascular, inflammatory or neurodegenerative diseases. Healthy subjects were volunteers with no known significant health issues. All data were acquired after obtaining prior written informed consent from the participants.

### 2.2 Neuropsychological Assessment

All participants underwent the same neuropsychological examination (at least 3 months post-stroke for patients) which included: the Free and Cued Selective Reminding test [32] (verbal anterograde memory); the DMS48 task [33] (visual anterograde recognition memory); the Stroop test [34] (inhibition); literal and semantic fluencies [34] (executive functions); D2 [35] (attention); digit-symbol test [36] (working memory); ExaDé confrontation naming test [37] (language) and three mood and affective scales: the State-Trait Anxiety Inventory [38], Starkstein Apathy Scale [39] and Beck Depression Inventory Scale [40].

### 2.3 MRI Acquisition

For the first 20 patients and 20 healthy subjects, 3D T1-MPRAGE sequences were acquired on a 3T scanner (Philips Achieva) with the following parameters: 1*1*1 mm voxel size, TE= 8.1ms, TR= 3.7ms, flip angle= 8°, FOV= 240*240, slice number= 170. 3D T2-FLAIR (Fluid Attenuated Inversion Recovery) sequences parameters were: 1*1*1 mm voxel size, TE= 337ms, TR= 8000ms, TI= 2400ms, FOV= 240*240, slice thickness= 1 mm, slice number= 170. For the next 20 patients and 25 healthy subjects, 3D T1-MPRAGE sequences were acquired on a 3T scanner (Philips Achieva) with the following parameters: 0.9*0.9*0.9mm voxel size, TE= 8.1ms, TR= 3.7ms, flip angle= 8°, FOV= 256x256, slice number= 189. 3D T2-FLAIR sequences parameters were: 1*1*1 mm voxel size, TE= 343ms, TR= 8000ms, TI= 2400ms, FOV= 240*240, slice thickness= 1 mm, slice number= 160.

### 2.4 IA and IA Variants Identification Protocol

Two raters (JV, KR) independently assessed the presence or absence of IA, its anatomical variants and if the lesion involved IA among patients. A standardized protocol was set up on MRIcron [41] with a standardized zoom (3) and contrast value (2000) adapted to the MRI sequence type (Fig. 1). First, the reviewer examined the axial slices from the bottom to the top. An IA was deemed present if a structure connecting both thalami was observed on at least one slice between the anterior and the posterior commissures. This presence had to be confirmed on coronal and sagittal slices and in case of doubt, the FLAIR sequences were used. In the event of kissing thalami, partial volume, or discordance between the two raters after attempts to reach a consensus, the subject was excluded from analyses. An IA was identified as damaged if at least one voxel of the lesion extended into it.

**Figure 1:**
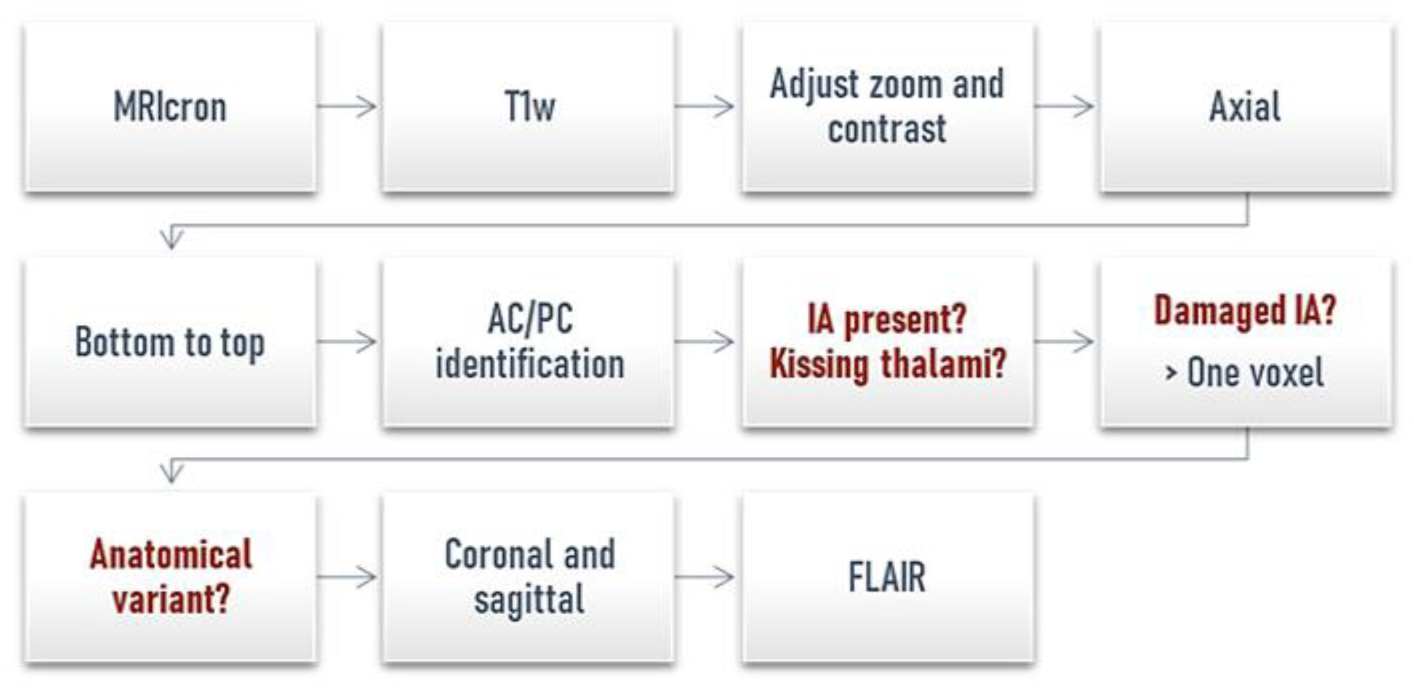
Standardized protocol to study IA. This protocol aims to identify the prevalence of IA, an eventual thalamic lesion extending into it and to characterize its anatomical form by two different raters using MRIcron software.

Regarding anatomical variants of IA, we followed the suggestions provided by Tsutsumi et al., 2021 [8]. Given there were two raters, we more specifically characterized IA as a broad form when it adhered to at least 1/3 of both thalami lengths on axial slices or 1/3 of their heights on coronal slices. The two raters did not have difficulty reaching agreement except in one case of possible bilobar IA.

### 2.5 Lesion Location

Lesions were manually segmented on the native T1w images by two independent investigators (JV, LD) using MRIcron software. Both native images and their corresponding lesions were normalized on the Montreal Neurological Institute (MNI) template and localized using the digitized Morel’s atlas [42]. The volume (mm^3^) of the normalized lesion in each nucleus and mammillothalamic tract per patient was then computed. Nuclei were gathered into nuclear groups using Morel’s repartition [43]. If a lesion reached the mammillothalamic tract (MTT) with a volume greater than 5mm^3^, the MTT was considered to be disrupted. For visualization purposes, all lesions were overlapped on the MNI152 template using MRIcroGL.

### 2.6 Statistical Analyses

#### 2.6.1 Bayesian Analyses

We employed Bayesian analyses using JASP [44]. The Bayesian paradigm stands apart from the frequentist paradigm as it does not rely on a cut-off for accepting or rejecting alternative hypotheses. Instead, it quantifies the strength of evidence in favor of a particular model using a continuous measure known as the Bayes factor. This Bayesian approach also permits the demonstration of evidence supporting the null model (BF_10_ < 0.3) as much as the absence of evidence when data are equally well predicted under both models (BF_10_ = 1). No p values are used in this context. BF_10_ are usually interpreted as follows: < 0.3: moderate evidence for the null model, 0.3-1: anecdotal evidence for the null model, 1: no evidence, 1-3: anecdotal evidence for the alternative model, 3-10: moderate evidence for the alternative model, 10-30: strong evidence for the alternative model, 30-100: very strong evidence for the alternative model, > 100: extreme evidence for the alternative model.

#### 2.6.2 Neuroimaging Analysis

For neuroimaging analysis, we compared the extent of lesions at different locations between groups using a Bayesian within-between subjects rmANOVA on the mean lesion volume per nuclear group among patients without or with an intact or damaged IA.

#### 2.6.3 Prevalence Analysis

To conduct analyses of IA presence or mammillothalamic tract disruption and also to compare laterality of thalamic infarct between groups, we used a Bayesian multinomial test. For analysis of mean age and mean education depending on the presence of an IA, we employed a Bayesian t-test. Where there were more than two groups to compare, a Bayesian ANOVA was utilized. If the Bayesian ANOVA yielded evidence toward the alternative hypothesis (BF_10_ > 3), we performed post-hoc t-tests to weigh up each pair of means.

#### 2.6.4 Neuropsychological Analyses

The scores obtained from each neuropsychological test were standardized to z-scores using normative scales. The psycho-affective scales were analyzed using raw data due to a lack of adequate normative scales. To compare subjects with and without an IA, we conducted a within-between subjects repeated measures ANOVA (rmANOVA) using z-scores from the FCSRT 3 total recall, digit-symbol, literal and semantic fluencies, Stroop interferences minus denomination (response time), DMS48 set 2 (response time to a forced choice recognition at a 1-hour delay), D2 (rhythm, GZ-F, number of processed items minus errors) and the confrontation naming test. These subtests were selected to minimize multiple comparisons and selection was based on literature-driven assumptions. To confirm it was possible to use the rmANOVA, we assessed normality using Shapiro-Wilk’s test, homogeneity of variance using Levene’s tests and visually inspected the data with QQ-plots. In cases of large violation of normality assumptions and for the study of psychoaffective scales, we resorted to Bayesian Mann-Whitney tests to weigh up the two groups of healthy subjects. To compare the three groups of patients and the healthy subject group and due to a lack of a Bayesian non-parametric alternative to the ANOVA, frequentist Kruskall-Wallis tests followed by Bonferroni-corrected Dunn’s tests were utilized. In the event of a between-subject effect (grouping effect) in the rmANOVA (BF_10_ > 3), we employed a Bayesian post-hoc t-test to identify differences between groups in the neuropsychological subtests of interest since a post-hoc function comparing groups depending on factors has not yet been implemented in the software. The posterior distributions of performance across tests by group were reconstructed to identify overall distinctions between patients and healthy subjects. Subsequently, we displayed the average z-score for each neuropsychological test within each group, aiming to identify the most discriminating neuropsychological tests from those selected for the rmANOVA. Finally, we used a Bayesian ANOVA followed by a Bayesian post-hoc t-test to quantify which test was the most discriminative one and represented significant results as boxplots using z-scores by groups to analyze individual performances.

## 3. Results

### 3.1 Neuroimaging Analyses

Two cases of kissing thalami were identified among healthy subjects and led to their exclusion (MRI in Supp. Fig. 1). The initial concordance between the two independent raters about the presence or absence of IA was 95% (kappa=0.89) and 99% after attempts to reach a consensus which led to the exclusion of one healthy subject. The analyses finally included 42 healthy subjects and 40 patients. There were no differences between the two groups in terms of age, education and gender (Tab.1).

**Table 1:** Demographical Data of All Subjects.

|  | **Healthy Subjects (n=42)** |  | **Patients (n=40)** |  |  |
| --- | --- | --- | --- | --- | --- |
|  | **With IA** | **Without IA** | **Intact IA** | **Damaged IA** | **Absent IA** |
| *N (women)* | 32 (20) | 10 (3) | 18 (19) | 12 (6) | 10 (0) |
| *Mean age* | 47.9 | 54.1 | 48.4 | 54.8 | 51.6 |
| *(SD, min, max)* | (15.2, 20, 69) | (5.9, 42, 62) | (13.2, 23, 69) | (18.9, 24, 75) | (13, 25, 74) |
| *Mean years of study* | 13.4 | 13.8 | 12.1 | 12.8 | 12.5 |
| *(SD, min, max)* | (3.7, 5, 21) | (3.1, 9, 17) | (3.2, 5, 17) | (4.3, 4, 17) | (2.7, 9, 17) |

A 3D representation of thalamic nuclei from a healthy subject, including IA, is represented in Figure 2. Please note how close the mammillothalamic tract (in pink) is to the IA.

**Figure 2:**
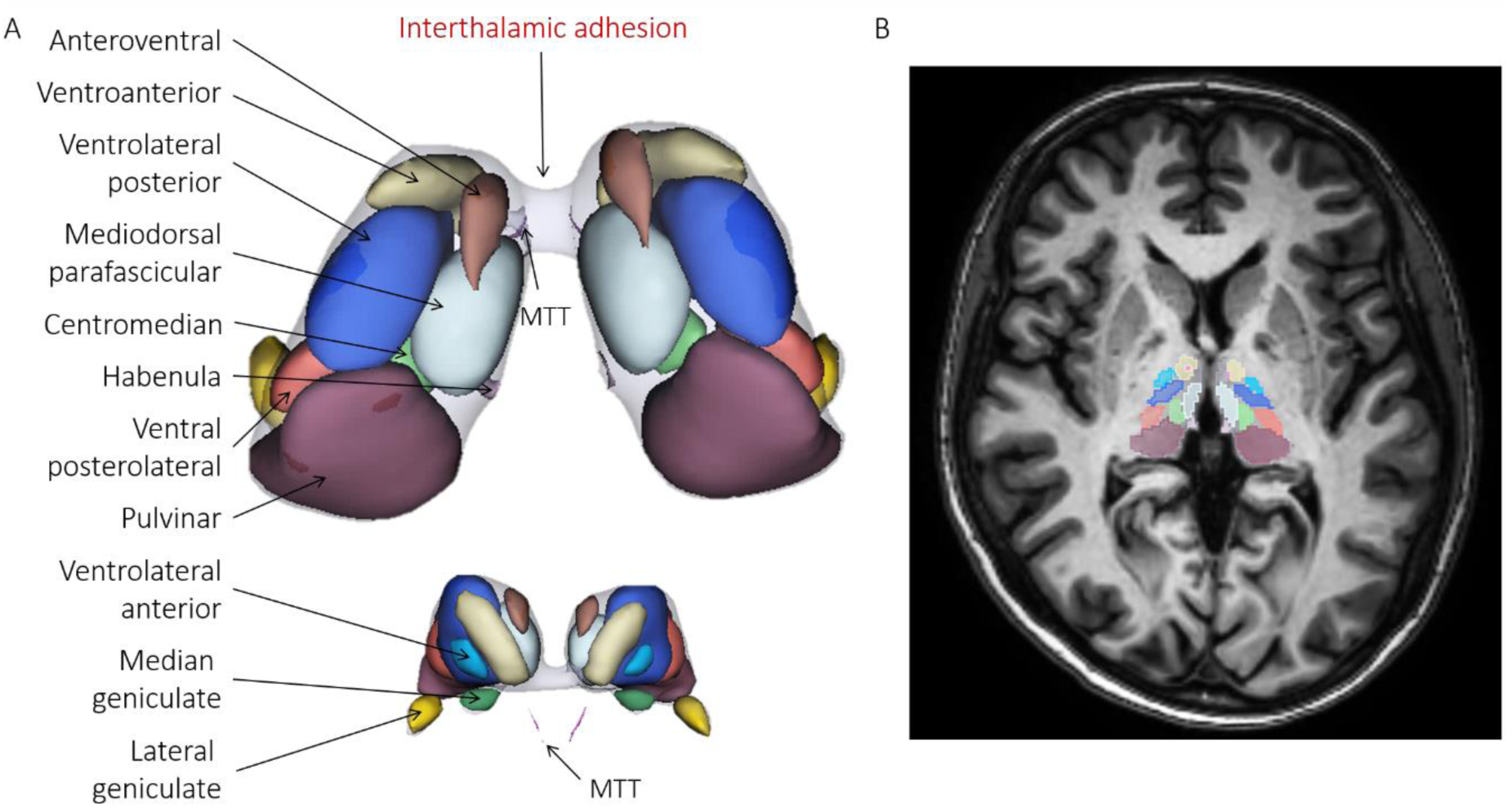
(A) 3D representations of thalamic nuclei of a single healthy subject in axial (left) and (B) corresponding axial MRI slice. MTT: Mammillothalamic Tract. Segmentations were generated using T1w images in Thalamus Optimized Multi-Atlas Segmentation (THOMAS) using the HIPS method [45, 46].

The prevalence of IA and its anatomical variants are presented in Table 2 (see also Fig. 3). IA was absent in 24% of all participants. IA absence was more frequent among men (39%) than women (8%) (BF_10_ independent multinomial test = 51). We found moderate evidence favoring the absence of an age effect (BF_10_ t-test = 0.38) or education (BF_10_ t-test = 0.27). In addition, no impact of age (BF_10_ ANOVA= 0.19, education (BF_10_ ANOVA = 0.16) or gender (BF_10_ independent multinomial = 0.29) on the distribution of the variants was evidenced, neither across all subjects nor within healthy subjects or patient subgroups. Prevalence of IA did not differ between the groups of patients and healthy subjects (BF_10_ independent multinomial = 0.23).

**Figure 3:**
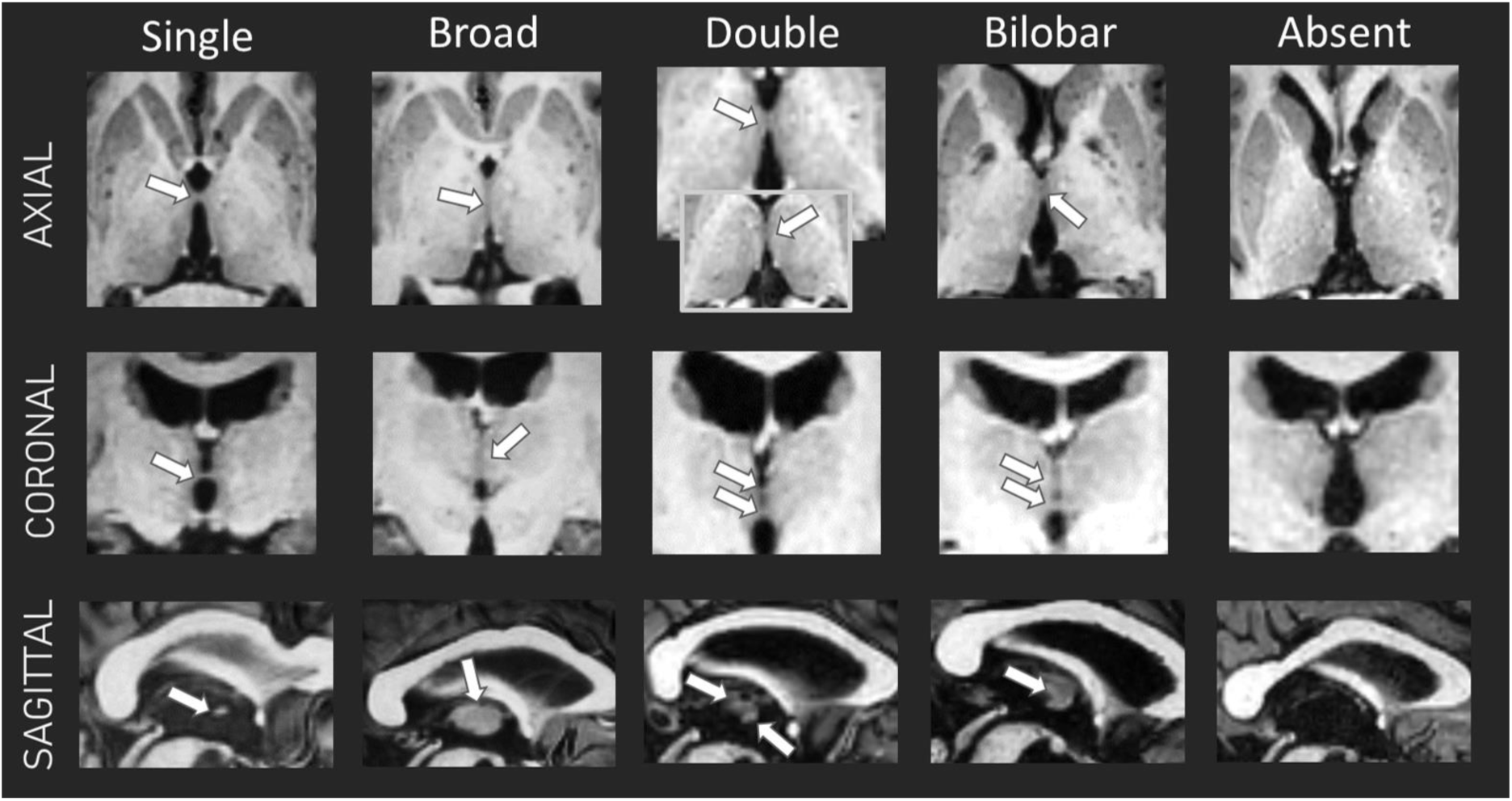
Illustration of the different anatomical variants of IA or illustration of its absence in single healthy subjects. White arrows indicate IA location.

An IA-related lesion was identified among 12 patients (Fig. 4). For those with a double IA, two patients had one damaged IA while other one was preserved (Supp. Fig. 2).

**Figure 4:**
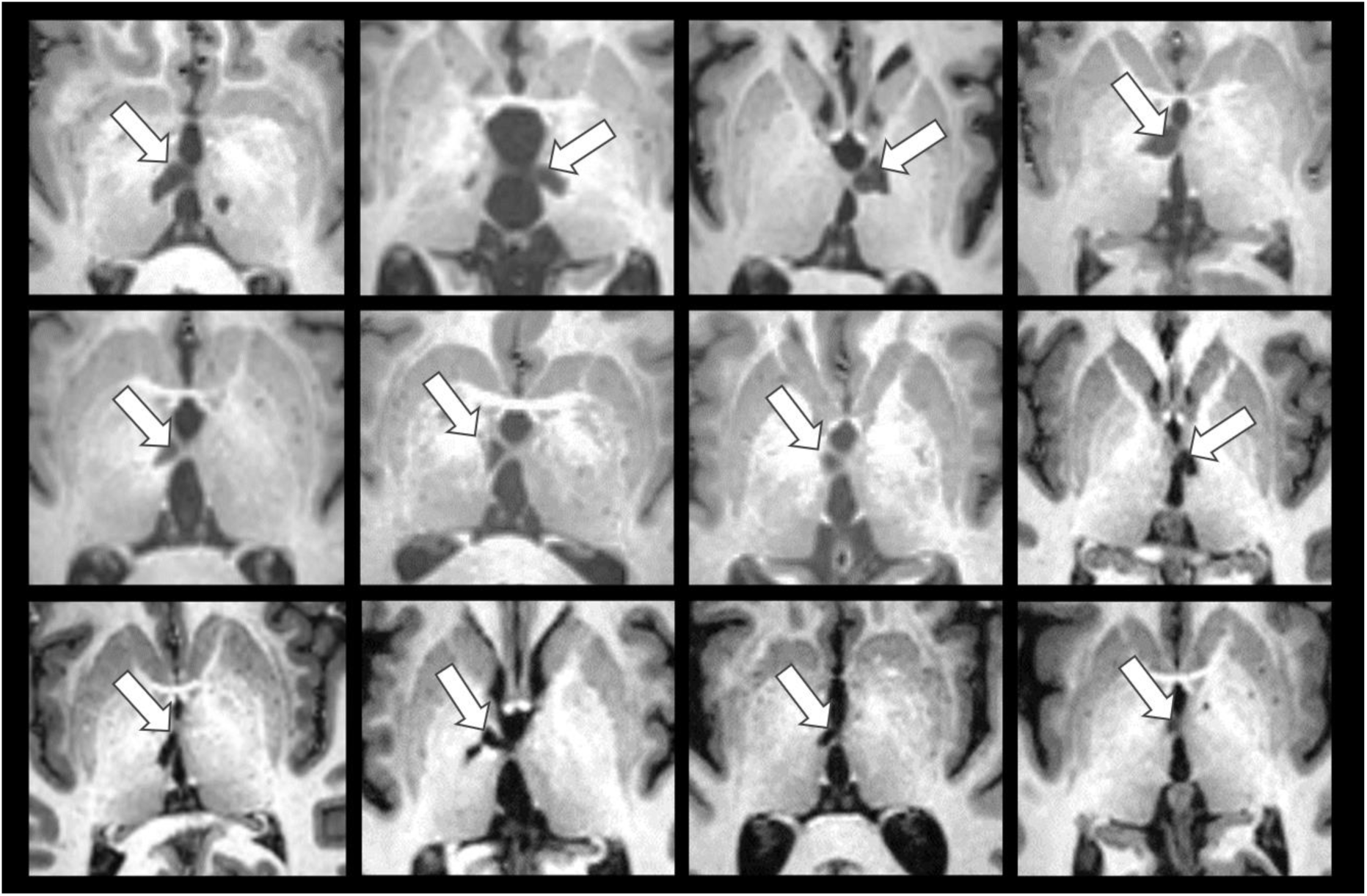
T1w axial slice for each of the 12 patients with a thalamic lesion reaching the IA (white arrows).

The mean lesion volume differed between the groups of patients (BF_10_ ANOVA = 6.7; IA intact: 285 ± 258 mm3; IA damaged: 604 ± 323 mm^3^; IA absent: 327± 201 mm^3^) and was higher among patients with a damaged IA than in patients with an intact IA (post-hoc BF_10_ t-test= 8). However, laterality of infarcts (BF_10_ independent multinomial test = 0.85) and the number of patients with a disrupted mammillothalamic tract (intact IA: 2; damaged IA: 5; IA absent: 3; BF_10_ independent multinomial = 0.88) were not dissimilar between the 3 subgroups of patients. Locations of infarcts were also distributed identically using the mean lesioned volume per nuclear group (BF_10_ rmANOVA = 0.31). Those lesions were mainly located in the median and lateral left thalamus for the 3 subgroups of patients (normalized lesions overlapping on an MNI template in Supp. Fig. 3). There was no significant difference between patient subgroups in terms of mean age (BF_10_ ANOVA = 0.29) or years of education (BF_10_ ANOVA = 0.21).

### 3.2 Neuropsychological Analyses

#### 3.2.1 Healthy Subjects

No difference, and even moderate evidence in favor of the null hypothesis, was observed between healthy subjects *with* (n=32) and *without* (n=10) an IA on neuropsychological tests (BF_10 rm_ANOVA = 0.28 for the factor group, BF_10_ = 0.09 for the factor group*tests). As the confrontation naming test violated normality assumptions, significant difference was assessed, along with psychoaffective scales, using Bayesian Mann-Whitney tests. No discrepancy between subjects with and without an IA was found either as analyses provided anecdotal evidence supporting null hypothesis (BF_10_ Mann-Whitney confrontation naming = 0.42; Starkstein BF_10_ = 0.50, Beck BF_10_ = 0.35, Spielberg BF_10_=0.39).

**Table 2:** Frequency of the absence or presence of IA and its different anatomical variants among healthy subjects (n=42), patients (n=40) and all subjects (n=82). The prevalence of IA variants among patients was computed on 39 patients due to a discordance between the two raters.

|  | One | Broad | Double | Bilobar | Absent |
| --- | --- | --- | --- | --- | --- |
| **Healthy subjects** | 20 (48%) | 6 (14%) | 3 (7%) | 3 (7%) | 10 (24%) |
| **Patients** | 14 (36%) | 4 (10%) | 7 (18%) | 4 (10%) | 10 (25%) |
| **All subjects** | 34 (42%) | 10 (12%) | 10 (12%) | 7 (10%) | 20 (24%) |

The only IA variant that counted enough subjects to perform statistical analyses were the broad variant group (n=6). There was anecdotal evidence favoring the null hypothesis between the healthy subject group with a typical single anatomical variant (n=20) of the IA and those with a broad variant (n=6) (Bayesian rmANOVA: BF_10_=0.46 for the factor group effect, BF_10_=1.3 for group*test; Bayesian Mann-Whitney test applied to the confrontation naming test = BF_10_=0.46). As other anatomical variants were represented by groups with fewer than 5 subjects, no further statistical analyses were conducted.

Given the evidence of an absence of neuropsychological differences between healthy subjects with and without an IA, the two groups were unified into a single one (n=42) to improve the statistical power of further analyses. To ensure reliability of the results, all analyses presented hereafter were also performed using only the group of healthy subjects with an IA, which led to the same conclusions, albeit with decreased statistical power.

#### 3.2.2 Patients vs Healthy Subjects

The rmANOVA showed strong evidence of differences in terms of the performance between the patients and healthy groups (BF_10_=3.3×10^10^ for the factor group, BF_10_=1.2×10^10^ for the interaction group*test). These variations are represented in Figure 5A using posterior distributions. Mean z-scores per group per neuropsychological subtest used in the rmANOVA are depicted in Figure 5B. Subsequent post-hoc Bayesian t-tests on the group variable provided extreme evidence of discordance between the healthy subject group and each group of patients. Patients *with* an IA showed the lowest BF_10_ (157) while patients *without* an IA showed extreme evidence of a disparity with the highest BF_10_ (10648) and no overlap in the confidence intervals of this group and that of healthy subjects. In these conditions, BF10 values may be influenced by the number of subjects per group. However, this group of patients *without* an IA actually had the lowest number of subjects (10 vs 18 for an intact IA), suggesting that this effect did not drive the BF_10_ value. The group of patients with a damaged IA had a BF_10_ of 163. It should be noted that results were in favor of the null model (BF_10_ < 0.35) regarding dissimilarities between the three groups of patients.

**Figure 5:**
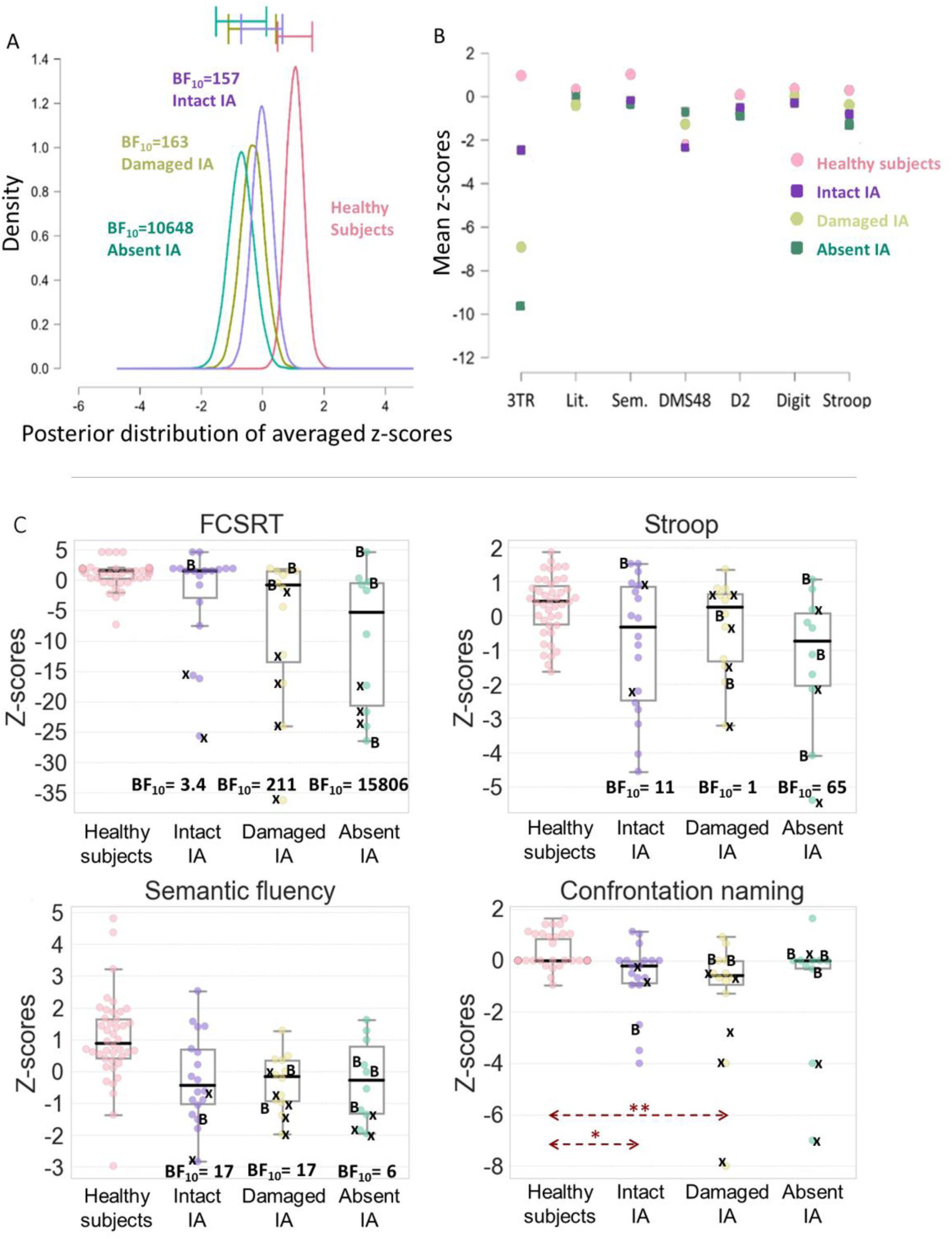
(A) Posterior distributions of the mean z-score per group to the neuropsychological tests used in the rmANOVA. BF_10_ results are from the post-hoc t-test on the group factor against the healthy subject group. Error bars represent 95% confidence intervals. (B) Mean z-scores per group per neuropsychological subtests used in the rmANOVA. FCSRT 3 Total Recall (3TR), literal fluencies (Lit.), semantic fluencies (Sem.), DMS48 (set 2), D2 (GZ-F), digit-symbol (Digit), Stroop (interferences minus denomination). (C) Boxplots of the FCSRT 3 Total Recall, Stroop (I-D), Semantic fluency and Confrontation naming test by groups. BF_10_ results are from an ANOVA followed by a post-hoc t-test. A black x indicates a lesion to the mammillothalamic tract and a “B” indicates a bilateral thalamic lesion. Groups: Healthy subjects (n=42), Intact IA (n=18), Damaged IA (n=12), Absent IA (n=10).

To further explore the most discriminative tests employed in the rmANOVA, we used Bayesian ANOVAs which revealed differences between the patient and healthy subject groups on the FCSRT (BF_10_ =253), Stroop test (BF_10_=14) and semantic fluencies (BF_10_ =129) as well as in the confrontation naming (Kruskal-Wallis test followed by a Dunn’s test with a Bonferroni correction). Boxplots depicting those results are presented in Figure 5C along with results from the post-hoc Bayesian t-test following the ANOVA.

The BF_10_ values (Fig. 5C) demonstrated that the group of patients *without* an IA (BF_10_ = 15806) exhibited a trend towards more severe impairment compared to healthy subjects than the group of patients *with* an IA (BF_10_ = 3.4) on the FCSRT. A similar tendency was found for the Stroop test. It may be possible that such results for the FCSRT are driven by concomitant lesions to the mammillothalamic tract (patients denoted with an x in Fig. 5C) which are known to be related to more severe cognitive impairment in verbal memory [44]. When the patients with mammillothalamic tract lesions were removed, the group of patients *without* an IA still showed strong evidence of a difference with the group of healthy subjects (BF_10_=19) whereas there was no longer any variation between the group of patients *with* an IA (BF_10_=0.53) and healthy subjects on the FCSRT. There was no evidence of a discrepancy between the two groups of patients with and without an IA however (BF_10_=0.8). We performed the same analysis while removing all patients with MTT lesions and all outliers (from all groups, i.e., subjects < or > to 1.5 inter-quartile interval). This did not change the results of this analysis.

There was evidence for the null model when comparing the four groups on the psychoaffective scales (Bayesian ANOVA: Starkstein BF_10_=0.17, Beck BF_10_=0.17, Spielberg BF_10_=0.5).

## 4. Discussion

In this study, IA was absent in 24% of the subjects. Women had a higher prevalence of IA (92%) compared to men (61%) while no effect of age was evidenced. These results are in line with the previous existing literature [4, 7, 10, 13, 14, 16, 17]. We also observed several IA variants from the most typical to the most unusual (Fig. 3) although these variants are usually not studied. In this context, IA appears an intriguing brain structure since the reasons for such an overall variability are unclear.

Another mystery relates to the functional role of IA. The literature is very scarce regarding any contribution to cognition which appears to contradict recent findings that the IA may be a white matter tract [7, 27]. In our study, the presence or absence of an IA had no significant effect on the performance on several neuropsychological tests in healthy subjects. An interpretation of these results suggests that IA plays no part or only has a marginal significance in cognition among the healthy population. This position can be supported by the idea that 20-25% of the population do not have an IA, implying that it is not necessary for any critical cognitive function. It is worth mentioning however that the role of several nuclei in the thalamus such as the mediodorsal nucleus are in fact poorly understood [47]. Another interpretation is therefore that without a suitable theory about the role of IA, we did not use the appropriate tests to assess its contribution.

Given this situation, we aimed to test the role of IA in patients with isolated thalamic strokes. We thought that the presence of an IA could be a protective factor against cognitive impairment through post-stroke functional reorganization. In contrast, we hypothesized that patients *without* an IA would show the most severe impairment. The results of this study confirm these predictions as the group of patients *with* an IA was the least severely impaired compared to the group of healthy subjects while the group of patients *without* an IA was the performed the most poorly during the general analysis on all tests combined (Fig. 5A). The group of patients *with* an IA *and* a lesion extending in it showed an intermediary profile of cognitive impairment between the two other groups of patients.

Follow-up analyses on the neuropsychological tests showed that patient performance was not affected on all tests. This was expected as not all the tests are sensitive enough to be impaired by thalamic lesions. Tests that were more impacted were those with a strong verbal component (the verbal memory task FCSRT, semantic fluency and confrontation naming) in accordance with patient lesion placements which were mostly left-sided (28 left, 6 right, 6 bilateral) and the known role of the left thalamus in lexical selection [48]. Patients also produced poor results on the Stroop test which strongly depends on the frontal lobes. This also appears to tally with the overall pattern of patient lesions which mainly encompassed the mediodorsal but also ventrolateral thalamus [47].

The patients *without* an IA performed worse than the patients *with* an IA compared to the group of healthy subjects on both the verbal memory task and the Stroop test. Focusing on the verbal memory task, where the largest differences were observed, we obtained similar results (though with lower BF_10_) after removing outliers, or patients with mammillothalamic tract lesions, which are known to impair memory. Assessing the semantic fluency task, the reverse trend was observed (patients *with* an IA were more impaired than patients without IA compared to healthy subjects) but for much lower BF_10_ differences. Importantly, these effects could not be accounted for by age, laterality of the infarct, volume, or localization of the lesion (cf. 3.1 Neuroimaging Analyses). In addition, we did not find evidence of disparities between the groups in terms of anxiety or depression that could explain the results.

The case of thalamic lesions extending into the IA is more ambiguous. A similar trend of deficits than patients *without* an IA was observed in patients with a lesioned IA but could be due to larger lesions or a higher prevalence of concomitant lesions to the mammillothalamic tract. Additionally, it is possible that complete disruption of IA is necessary to induce deficits as a partial lesion may preserve some fibers and allow potential compensation by the opposite thalamus. Moreover, variations in IA anatomy may affect connectivity: a broad variant may support robust connectivity while standard and double variants may have thinner connections, limiting the number of crossing fibers [29].

The absence of an IA could thus be a prognosis of poor neuropsychological outcomes following thalamic strokes. These findings also lead to the hypothesis that IA may play a role in compensatory mechanisms rather than having a specific part in cognition. In the absence of IA, the cognitive circuits in healthy subjects otherwise depending on this structure may be supported by other midline structures such as the anterior and posterior commissures or the corpus callosum, potentially explaining the absence of neuropsychological differences between healthy subjects with and without an IA. However, in the case of a thalamic lesion, IA-related circuitry may become important to support compensatory mechanisms, resulting in the lower neuropsychological deficits observed in patients with preserved IA. If confirmed, accurate identification of IA could be crucial. It would become especially important for microsurgical and endoscopic approaches to the third ventricle and pineal region. For instance, in the microsurgical resection of tumors in the third ventricle, preserving the IA could minimize the risk of neuropsychological deficits.

Overall, these results require further exploration and extensive research on patients with isolated thalamic strokes. A limitation of our finding is that we did not actually find evidence in favor of differences between the groups of patients using the Bayesian analyses we conducted. This can be interpreted as resulting from a lack of power since the number of patients without an IA was rather low (n=10 patients). In addition, the variability of such neuropsychological investigations, due to various factors even if we tried to control them, such as age, educational level, concomitant mammillothalamic tract lesions, IA anatomical variants, exact location of lesions and volume of the lesion, increase the levels of variance that can only be fulfilled by larger groups of subjects.

In general, the wide variation observed in the literature regarding the size, prevalence, function and connectivity of IA could be attributed to the lack of standardization in MRI procedures. These discrepancies encompass differences in MRI acquisition parameters. For instance, thicker slices, poor resolution especially in DTI images and thick inter-slice gaps are all factors that can result in missing a narrow IA and lead to a higher rate of IA absence [24]. Results about IA incidence using MRI can also contrast with postmortem findings because of the inherent tissue processing that could lead to IA rupture, artificially decreasing its prevalence [17, 24, 49]. The protocol designed for this study aims to establish an initial standardized MRI protocol for IA investigations, serving as a foundational framework for further enhancement.

## 5. Acknowledgments

We would like to thank Hélène Gros, Nathalie Vayssière and the MRI Research Center for their help in MRI acquisition.

## 7. Supplementary files

**Supplementary Figure 1:**
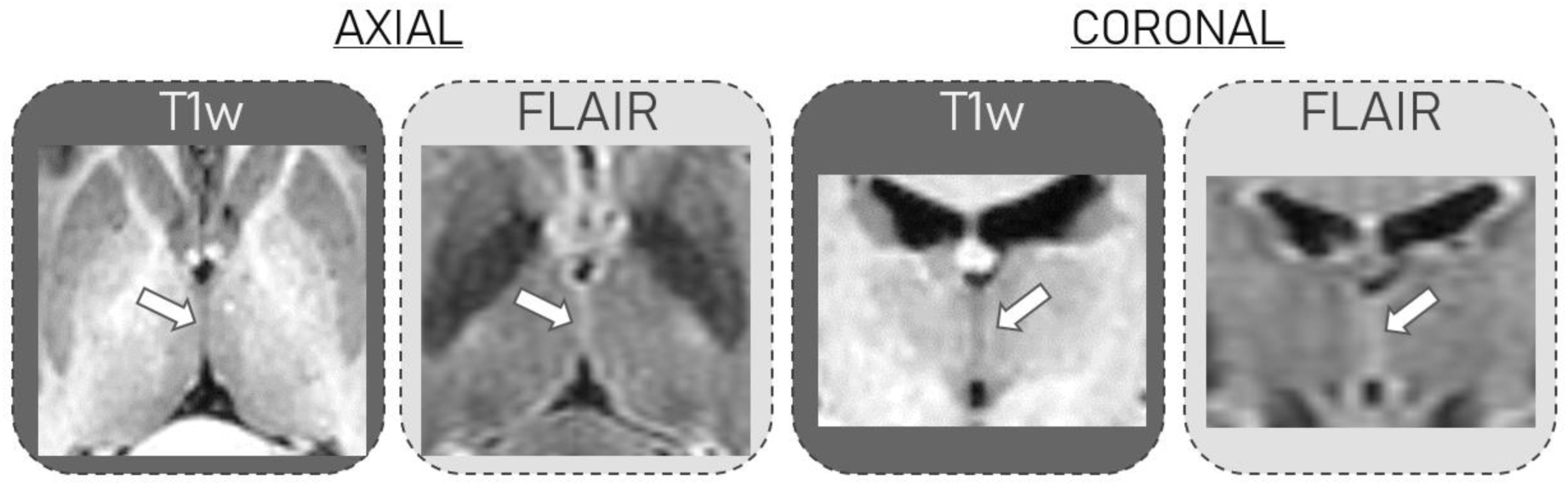
Illustration of a kissing thalami case on the axial and coronal slice of a T1w (dark gray) or a FLAIR (light gray) image which does not allow identification of the presence or absence of IA. Arrows indicate the contact between thalami where IA could have been identified if present.

**Supplementary Figure 2:**
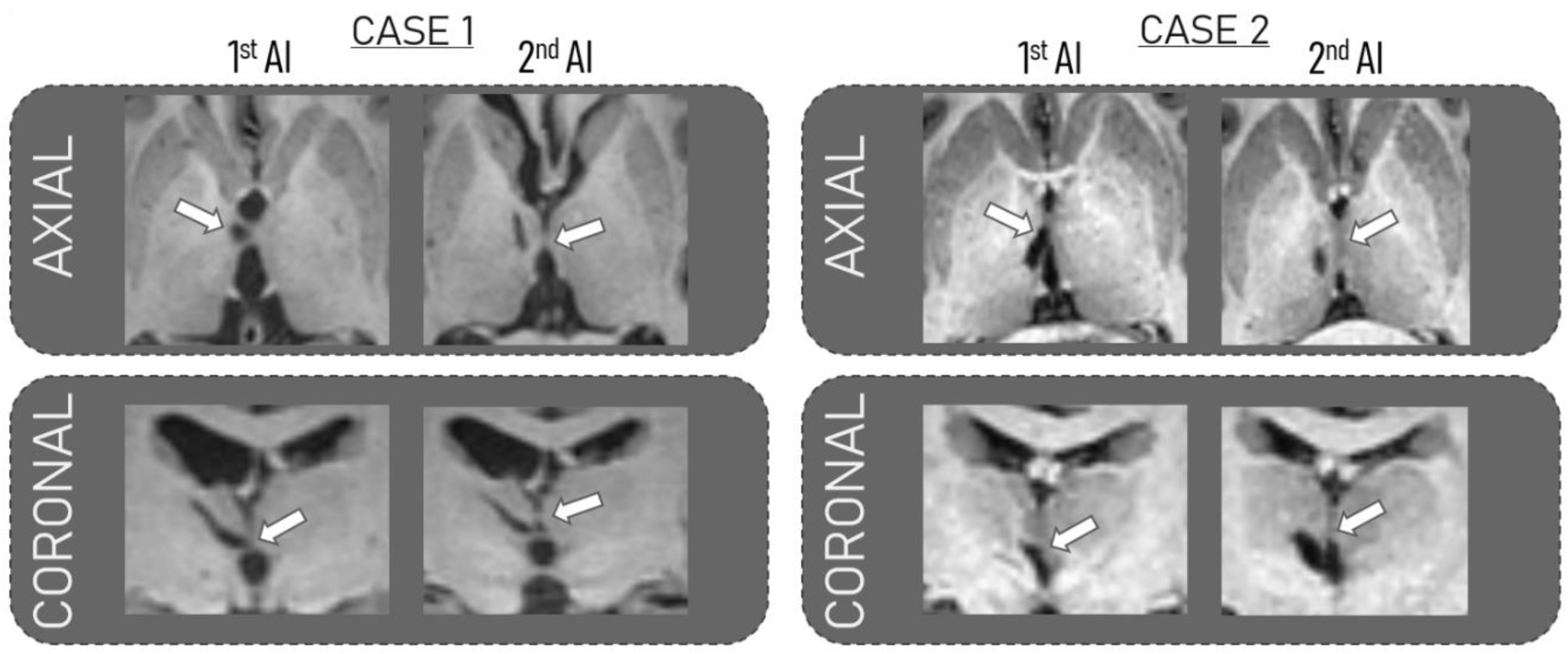
Illustration of an axial and coronal slice of a T1w image from two patients with a double IA. The first image indicates the lesioned IA while the second demonstrates it is preserved. White arrows indicate IA location.

**Supplementary Figure 3:**
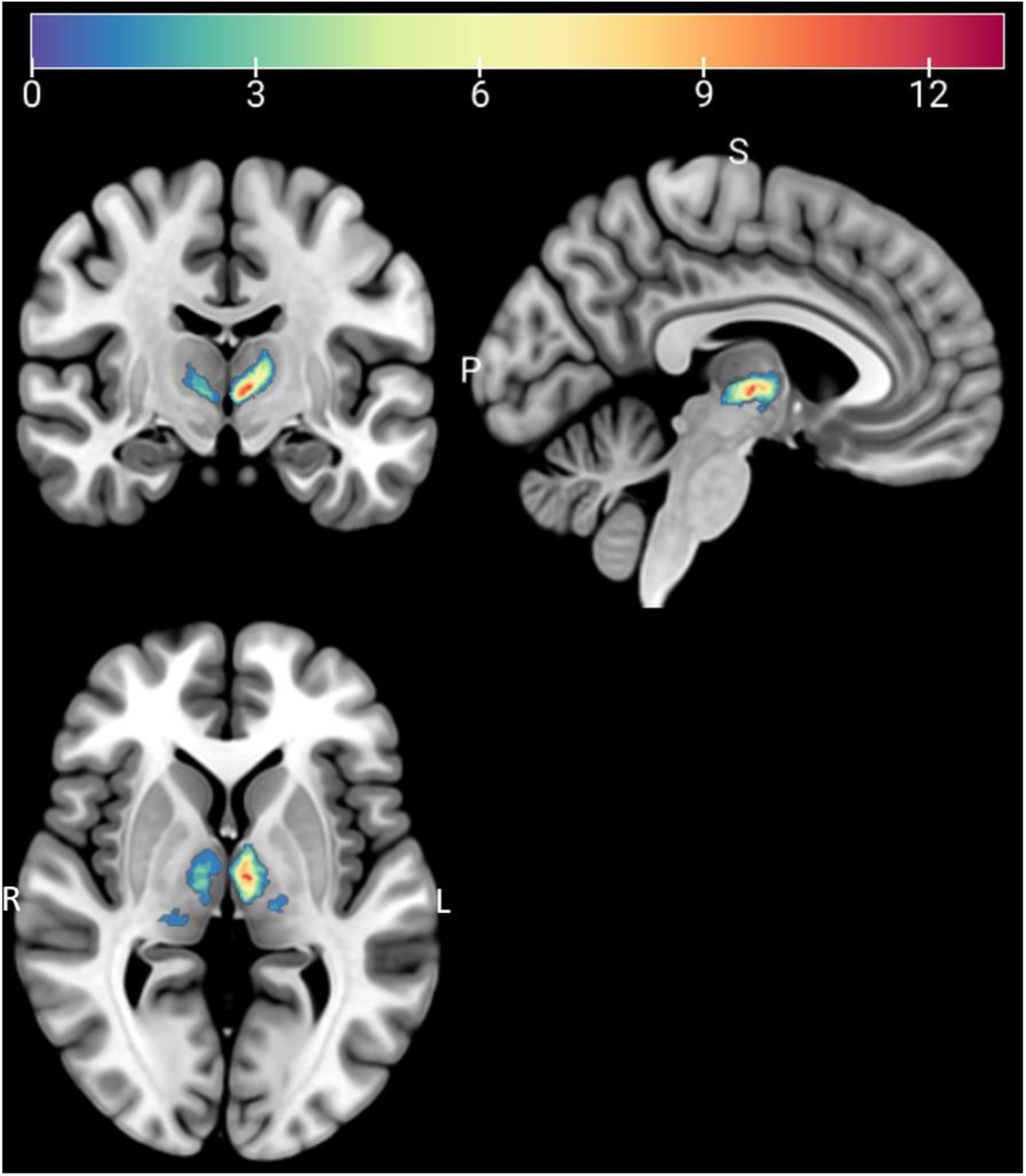
Representation of all lesions from the 40 patients on the MNI152 template after normalization. The scale bar represents the number of overlapping lesions. L: Left; R: Right; S: Superior; P: Posterior.

## References

[1] Patra, A., Ravi, K. S., & Asghar, A. (2022). The Prevalence, Location, and Dimensions Of Interthalamic Adhesions and Their Clinical Significance: Corpse Brain Analysis. Asian Journal of Neurosurgery, 17(04), 600–605.

[2] Pavlović, M. N., Jovanović, I. D., Ugrenović, S. Z., Kostić, A. V., Kundalić, B. K., Stojanović, V. R., … & Antić, M. M. (2020). Position and size of massa intermedia in Serbian brains. Folia Morphologica, 79(1), 21–27.

[3] Davie, J. C., & Baldwin, M. (1967). Radiographic-anatomical study of the massa intermedia. Journal of neurosurgery, 26(5), 483–487.

[4] Malobabic, S., Puskas, L., & Blagotic, M. (1987). Size and position of the human adhaesio interthalamica. Gegenbaurs Morphol Jahrb, 133(1), 175–180.

[5] Etus, V., Guler, T. M., & Karabagli, H. (2017). Third ventricle floor variations and abnormalities in myelomeningocele-associated hydrocephalus: our experience with 455 endoscopic third ventriculostomy procedures. Turk Neurosurg, 27(5), 768–771.

[6] Miller, E., Widjaja, E., Blaser, S., Dennis, M., & Raybaud, C. (2008). The old and the new: supratentorial MR findings in Chiari II malformation. Child’s Nervous System, 24, 563–575.

[7] Şahin, M. H., Güngör, A., Demirtaş, O. K., Postuk, Ç., Fırat, Z., Ekinci, G., … & Türe, U. (2023). Microsurgical and fiber tract anatomy of the interthalamic adhesion. Journal of Neurosurgery, 1(aop), 1–10.

[8] Tsutsumi, S., Ono, H., & Ishii, H. (2021). Massa intermedia of the thalamus: an anatomical study using magnetic resonance imaging. Surgical and Radiologic Anatomy, 43(12), 1927–1932.

[9] Viller, F. M. R. (1887). Recherches anatomiques sur la commissure grise (Doctoral dissertation, Verlag nicht ermittelbar).

[10] Borghei, A., Cothran, T., Brahimaj, B., & Sani, S. (2020). Role of massa intermedia in human neurocognitive processing. Brain Structure and Function, 225, 985–993.

[11] Borghei, A., Kapucu, I., Dawe, R., Kocak, M., & Sani, S. (2021). Structural connectivity of the human massa intermedia: A probabilistic tractography study. Human Brain Mapping, 42(6), 1794–1804.

[12] Rabl, R. (1958). Strukturstudien an der Massa intermedia des Thalamus opticus. J. Hirnforsch., 4, 78–112.

[13] Takahashi, T., Yücel, M., Yung, A. R., Wood, S. J., Phillips, L. J., Berger, G. E., … & Pantelis, C. (2008). Adhesio interthalamica in individuals at high-risk for developing psychosis and patients with psychotic disorders. Progress in Neuro-Psychopharmacology and Biological Psychiatry, 32(7), 1708–1714.

[14] Wong, A. K., Wolfson, D. I., Borghei, A., & Sani, S. (2021). Prevalence of the interthalamic adhesion in the human brain: a review of literature. Brain Structure and Function, 226, 2481–2487.

[15] Borghei, A., Piracha, A., & Sani, S. (2021). Prevalence and anatomical characteristics of the human massa intermedia. Brain Structure and Function, 226, 471–480.

[16] Nopoulos, P. C., Rideout, D., Crespo-Facorro, B., & Andreasen, N. C. (2001). Sex differences in the absence of massa intermedia in patients with schizophrenia versus healthy controls. Schizophrenia research, 48(2-3), 177–185.

[17] Erbaǧcı, H., Yıldırım, H., Herken, H., & Gümüşburun, E. (2002). A magnetic resonance imaging study of the adhesio interthalamica in schizophrenia. Schizophrenia research, 55(1-2), 89–92.

[18] Trzesniak, C., Linares, I. M., Coimbra, É. R., Júnior, A. V., Velasco, T. R., Santos, A. C., … & Crippa, J. A. (2016). Adhesio interthalamica and cavum septum pellucidum in mesial temporal lobe epilepsy. Brain imaging and behavior, 10(3), 849–856.

[19] Birnbaum, R., Parodi, S., Donarini, G., Meccariello, G., Fulcheri, E., & Paladini, D. (2018). The third ventricle of the human fetal brain: Normative data and pathologic correlation. A 3D transvaginal neurosonography study. *Prenatal Diagnosis*, 38(9), 664-672.

[20] Takahashi, T., Chanen, A. M., Wood, S. J., Walterfang, M., Harding, I. H., Yücel, M., … & Pantelis, C. (2009a). Midline brain structures in teenagers with first-presentation borderline personality disorder. Progress in Neuro-Psychopharmacology and Biological Psychiatry, 33(5), 842–846.

[21] Rosales, R. K., Lemay, M. J., & Yakovley, P. I. (1968). The development and involution of massa intermedia with regard to age and sex. Journal of Neuropathology and Experimental Neurology, 27(1), 166–166.

[22] Takahashi, T., Malhi, G. S., Wood, S. J., Yücel, M., Walterfang, M., Nakamura, K., … & Pantelis, C. (2010). Midline brain abnormalities in established bipolar affective disorder. Journal of affective disorders, 122(3), 301–305.

[23] Takahashi, T., Yücel, M., Lorenzetti, V., Nakamura, K., Whittle, S., Walterfang, M., … & Allen, N. B. (2009b). Midline brain structures in patients with current and remitted major depression. Progress in Neuro-Psychopharmacology and Biological Psychiatry, 33(6), 1058–1063.

[24] Trzesniak, C., Kempton, M. J., Busatto, G. F., de Oliveira, I. R., Galvão-de Almeida, A., Kambeitz, J., … & Crippa, J. A. S. (2011). Adhesio interthalamica alterations in schizophrenia spectrum disorders: a systematic review and meta-analysis. Progress in Neuro-Psychopharmacology and Biological Psychiatry, 35(4), 877–886.

[25] Malobabić, S., Puškaš, L., & Vujašković, G. (1990). Golgi morphology of the neurons in frontal sections of human interthalamic adhesion. Acta anatomica, 139(3), 234–238.

[26] Parra, J. E. D., Ripoll, Á. P., & García, J. F. V. (2022). Interthalamic adhesion in humans: a gray commissure?. Anatomy & Cell Biology, 55(1), 109–112.

[27] Laslo, P., Slobodan, M., NELA, P., MILOŜ, M., Rade, P., & Tatjana, I. (2005). Specific circular organization of the neurons of human interthalamic adhesion and of periventricular thalamic region. International journal of neuroscience, 115(5), 669–679.

[28] Damle, N. R., Ikuta, T., John, M., Peters, B. D., DeRosse, P., Malhotra, A. K., & Szeszko, P. R. (2017). Relationship among interthalamic adhesion size, thalamic anatomy and neuropsychological functions in healthy volunteers. Brain Structure and Function, 222, 2183–2192.

[29] Zhou SY, Suzuki M, Hagino H, Takahashi T, Kawasaki Y, Nohara S, et al. Decreased volume and increased asymmetry of the anterior limb of the internal capsule in patients with schizophrenia. Biol Psychiatry. 2003; 54:427–436.

[30] Kochanski, R. B., Dawe, R., Kocak, M., & Sani, S. (2018). Identification of stria medullaris fibers in the massa intermedia using diffusion tensor imaging. World Neurosurgery, 112, e497–e504.

[31] Danet, L., Barbeau, E. J., Eustache, P., Planton, M., Raposo, N., Sibon, I., … & Pariente, J. (2015). Thalamic amnesia after infarct: the role of the mammillothalamic tract and mediodorsal nucleus. Neurology, 85(24), 2107–2115.

[32] Van der Linden, M., Adam, S., Agniel, A., Baisset-Mouly, C., Bardet, F., & Coyette, F. (2004). L’évaluation des troubles de la mémoire épisodique: fondements théoriques et méthodologiques. M. VAN DER LINDEN, L’évaluation des troubles de la mémoire, Marseille, Solal, 11-20.

[33] Barbeau, E., Didic, M., Tramoni, E., Felician, O., Joubert, S., Sontheimer, A., … & Poncet, M. (2004). Evaluation of visual recognition memory in MCI patients. Neurology, 62(8), 1317–1322.

[34] Godefroy, O. “GREFEX.(2008).” Fonctions exécutives et pathologies neurologiques et psychiatriques.

[35] Brickenkamp, R. & Zillmer, E. (1998). The d2 Test of Attention. Seattle, Washington: Hogrefe & Huber Publishers.

[36] Wechsler, D. (2001). Echelle clinique de mémoire de Wechsler (MEM-III). 3th ed. Paris: Centre de Psychologie Appliquée.

[37] Bachy-Langedock, N. (1989). Batterie d’examen des troubles de la dénomination (ExaDé). Brussels: Editions du Centre de Psychologie Appliquée.

[38] Spielberger, C. D., Gorsuch, R. L., Lushene, R., Vagg, P. R., & Jacobs, G. A. (1983). Manual for the State-trait anxiety inventory (form Y self-evaluation questionnaire) consulting psychologists press: Palo Alto. CA.

[39] Starkstein, S. E., & Leentjens, A. F. (2008). The nosological position of apathy in clinical practice*. Journal of Neurology*, Neurosurgery & Psychiatry, 79(10), 1088–1092.

[40] Beck, A. T., & Steer, R. A. (1993). Manual for the Beck Anxiety Inventory. San Antonio, TX: Psychological Corp.

[41] Rorden, C., & Brett, M. (2000). Stereotaxic display of brain lesions. Behavioural neurology, 12(4), 191–200.

[42] Krauth, A., Blanc, R., Poveda, A., Jeanmonod, D., Morel, A., & Székely, G. (2010). A mean three-dimensional atlas of the human thalamus: generation from multiple histological data. Neuroimage, 49(3), 2053–2062.

[43] Morel, A. (2007). Stereotactic atlas of the human thalamus and basal ganglia. CRC Press.

[44] JASP Team (2023). JASP (Version 0.17.1.0)

[45] Su, J. H., Thomas, F. T., Kasoff, W. S., Tourdias, T., Choi, E. Y., Rutt, B. K., & Saranathan, M. (2019). Thalamus Optimized Multi Atlas Segmentation (THOMAS): fast, fully automated segmentation of thalamic nuclei from structural MRI. Neuroimage, 194, 272–282.

[46] Vidal, J. P., Danet, L., Péran, P., Pariente, J., Cuadra, M. B., Zahr, N. M., … & Saranathan, M. (2023). Robust thalamic nuclei segmentation from T1-weighted MRI using polynomial intensity transformation. arXiv:2304.07167.

[47] Pergola, G., Danet, L., Pitel, A. L., Carlesimo, G. A., Segobin, S., Pariente, J., … & Barbeau, E. J. (2018). The regulatory role of the human mediodorsal thalamus. Trends in cognitive sciences, 22(11), 1011–1025.

[48] Crosson, B. (1984). Role of the dominant thalamus in language: a review. Psychological bulletin, 96(3), 491.

[49] de Souza Crippa, J. A., Zuardi, A. W., Busatto, G. F., Sanches, R. F., Santos, A. C., Araújo, D., … & McGuire, P. K. (2006). Cavum septum pellucidum and adhesio interthalamica in schizophrenia: an MRI study. European Psychiatry, 21(5), 291–299.

